# Large effect life-history genomic regions are associated with functional morphological traits in Atlantic salmon

**DOI:** 10.1101/2024.12.05.627014

**Authors:** Tutku Aykanat, Paul V. Debes, Shadi Jansouz, Lison Gueguen, Andrew H. House, Annukka Ruokolainen, Jaakko Erkinaro, Victoria L. Pritchard, Craig R. Primmer, Geir H. Bolstad

## Abstract

Understanding pleiotropic architectures of phenotypes is instrumental for identifying the functional basis of adaptive genetic variation in the wild. Life-history variation may have a morphological basis that mediates resource acquisition allocation pathways, but identifying the underlying genetic basis of such traits is challenging. Using Atlantic salmon juveniles reared in common garden conditions, we test if two life-history associated loci in Atlantic salmon, s*ix6* and *vgll3,* are also associated with functional morphological traits. These loci had previously shown to exhibit strong signals of adaptation and are highly correlated with sea age at maturity. We show that genetic variation at the *vgll3* locus is linked to variation in morphological traits that underlie swimming performance, along a trade-off axis between efficient cruising and maneuvering, while the genetic variation at the *six6* locus was linked to variation in body-head proportions suggesting the potential functional importance of these traits for resource acquisition efficiency. However, the direction of changes in morphological traits associated with late-vs. early-maturing alleles was not always consistent with the expected direction of an effect to maturation timing. Our results reveal a complex morphological landscape associated with the genetic variation in these loci, possibly as a result of pleiotropy or linkage across these genomic regions.

## Introduction

Elucidating the genotype-phenotype relationship is instrumental in identifying the functional basis of adaptation and the potential targets of selection (Barrett and Hoekstra, 2011; Kawecki and Ebert, 2004; Nielsen *et al*, 2009; Wagner and Altenberg, 1996). Methodological and statistical advances in genetics now allow the identification of genomic regions responding to local selection, but the phenotypic link between genotype and fitness is often missing. Furthermore, many genomic regions can affect a multitude of phenotypic traits (via pleiotropy or linkage disequilibrium), which makes identifying the phenotypic traits under selection even more challenging (Paaby and Rockman, 2013; Solovieff *et al*, 2013). Even large, genome-wide association studies might be underpowered to detect pleiotropy (Wagner and Zhang, 2011), a methodological bias by which the adaptive significance of pleiotropy can be overlooked especially in natural, non-model populations even if it may have substantial effects to limit or promote adaptation (Orr, 2000; Otto, 2004). In natural systems, pleiotropy has been mostly studied at the phenotypic level to understand the evolutionary trade-offs by quantifying the physiological antagonisms between senescence and reproduction age. Yet, some studies identify pleiotropic effects of genes associated with adaptive genetic variation. For example, adaptive divergence between lake vs. marine morphs of threespine stickleback (*Gasterosteus aculeatus*) was explained by both pleiotropy in the *Eda* gene, and via strong linkage with the surrounding loci (Archambeault *et al*, 2020), illustrating the significance of such genetic architectures for adaptation.

Genomic regions with large adaptive importance are more likely to be detected in genome scans and are likely to have broad cellular and phenotypic effects (Shapiro *et al*, 2003, Jones *et al*, 2012, Barson *et al*, 2015). Major effect loci may also have broad cellular and phenotypic effects, with key regulatory roles across development, and could thereby drive strong genetic interdependences among traits (Chan *et al*, 2010; Colosimo *et al*, 2005; Jimenez-Gomez *et al*, 2010; Weith *et al*, 2023) but see (Boyle *et al*, 2017; Jakobson and Jarosz, 2020).

Therefore, loci exhibiting strong signals of adaptation may provide feasible empirical approaches to understand the significance of complex evolutionary processes, such as how pleiotropy and genetic interactions may hinder or promote adaptation in relation to the direction of natural selection between co-varying traits. Yet, such examples remain rare in non-model, wild organisms (Archambeault *et al*, 2020; Endler, 2000; Mackay *et al*, 2009; Olson-Manning *et al*, 2012).

Functional variation in morphological traits is integral in shaping resource use, and is correspondingly associated with life-history variation and ecotypic divergence in a large array of fish species, including salmonid fishes (Ferry-Graham *et al*, 2002; Fitzgerald *et al*, 2017; Keeley *et al*, 2005; Lundsgaard-Hansen *et al*, 2013; Mindel *et al*, 2016; Proulx and Magnan, 2004; Svanbäck and Eklöv, 2003; Taylor and McPhail, 1985; Varian and Nichols, 2010; Webb, 1988). For example, a fusiform body and shorter fins appear adaptive to a long-distance migratory strategy, likely because this body shape reduces the drag (hence the energy consumed) during migration (Morinville and Rasmussen, 2007). Likewise, a robust body shape with a larger head, caudal peduncle, and paired fins has been associated with piscivory in rainbow trout (*Oncorhynchus mykiss*, Keeley *et al*, 2005). Recent population genomics studies in Atlantic salmon have consistently identified two unlinked genomic regions, surrounding the genes *six6* in chromosome 9 and *vgll3* in chromosome 25 as strong targets for local selection within all European Atlantic salmon lineages (Barson *et al*, 2015; Miettinen *et al*, 2023; Pritchard *et al*, 2018; Zueva *et al*, 2020). These two loci have also been linked to a suite of life-history traits that strongly influence fitness. In particular, the *vgll3* and *six6* loci are strongly associated with sea age at first maturity (age at maturity from here on) in seagoing salmon at the individual– and population-levels respectively (Barson *et al*, 2015, see also, Ayllon *et al*, 2015; Sinclair-Waters *et al*, 2020), and the *vgll3* locus affects other alternative life-history strategies, such as precocious male maturation (Debes *et al*, 2021), and iteroparity (Aykanat *et al*, 2019). For example, individuals with late maturation genotype in the *vgll3* gene (*vgll3*^LL^) tended to stay at sea, on average, more than one year more than individuals with early maturation (*vgll3*^EE^) genotype (Barson *et al*, 2015, Mobley *et al*, 2021). This pattern was also pertinent during the freshwater phase within which juvenile males with *vgll3*^LL^ genotype matured at a much higher probability (∼ 60%) than juvenile males with *vgll3*^LL^ genotype (Debes *et al*, 2021).

Similarly, these loci explain variation across a number of metabolic and behavior traits. For example, *vgll3*^E^ allele has been shown to be associated with reduced aggression, increased maximum metabolic rate and aerobic scope at the juvenile stage (Bangura *et al*, 2022; Prokkola *et al*, 2022), while *six6* locus predicts diet acquisition patterns at sea in maturing adults with *six6*^E^ allele linked to increased stomach weight despite decreased foraging efforts (Aykanat *et al*, 2020). Likewise, *six6* locus exhibits epistasis with *vgll*3 to affect aerobic scope variation (Prokkola *et al*, 2022). The strong genetic interdependence of these phenotypic traits across two loci points to the potential for a shared developmental origin (of phenotypes to the respective loci), which is supported by the role of *vgll3* and *six6* in the hormonal and cell-fate commitment gene-expression pathways (Ahi *et al*, 2022; Kurko *et al*, 2020; Verta *et al*, 2020; Verta *et al*, 2024) and broad expression of *six6* during early development (Moustakas-Verho *et al*, 2020). Finally, a recent study showed both loci are linked to early life survival in juvenile Atlantic salmon, albeit via parental genetic effects, further supporting broad functional roles across life stages (Aykanat *et al*, 2024). This collective evidence indicates that these loci are broad-scale modulators of salmon phenotype. While many of these genetic effects have been demonstrated for traits at the juvenile stage, including early male maturation, metabolism, and behavior, any concordance with morphological variation, if any, is not known, but may be important to identify the architecture of personality traits, underlying genetic basis, and evolutionary constrains (Bangura *et al*, 2022, Kern et al, 2016).

To characterize the morphological variation associated with these locally adaptive genomic regions, we collected morphometric data form more than 700 juvenile immature Atlantic salmon at time when male parr maturation may take place (Debes *et al*, 2021). The fish was reared in common garden conditions and the association between morphology, and *six6* and *vgll3* genotype was estimated with a mixed-effect model that also controls for confounding length and condition factor effects. We identified a suite of morphological characters correlated to the genetic variation in both loci. Our study extends the broad pleiotropic effect of *vgll3* and *six6* loci to functional morphological traits, including phenotypes that are likely important targets of natural selection.

## Materials and Methods

### Fish breeding and, rearing and common garden conditions

Details of the breeding and rearing of individuals from the same cohort were provided in Debes *et al* (2021), using a similar experimental procedure and facilities, except that the fish in the present study were reared in approximately 2°C cooler water temperature (see, Debes *et al* (2019) for the details of the rearing temperature). Briefly, the parents of the individuals used were from Baltic lineage Atlantic salmon from a broodstock maintained by the Natural Resources Institute Finland (Laukaa, Finland) that were initially originated from the Neva River in Russia (Gulf of Finland in the Eastern Baltic Sea) and has been used for juvenile salmon supplementation in the Kymi river in South-Eastern Finland (Asheim *et al*, 2023). Parents were crossed using a biallelic cross design where 44 parents were crossed in a 11 two by two factorials design, in which, each factorial unit composed of a pair of males and females which are homozygote in early (E) and late (L) maturing alleles in the *vgll3* locus (i.e. *vgll3*^EE^ and *vgll3*^LL^ genotypes, Supplementary Table 1), with one type of genotype within each of the full-sib families with EE, EL, LE, and LL genotypes in the *vgll3* locus (Supplementary Table 1). *Six6* genotype distribution was not considered during the breeding design, but was nevertheless segregating in a number of families (Supplementary Table 1). Crosses were performed in November 2017, and offspring were reared in the University of Helsinki fish facilities, first in replicated family-specific compartments in vertical incubators until the swim-up stage. Subsequently offspring of all families were mixed in eight tanks with roughly balanced proportions of families per tank (Supplementary Table 2). In August 2018, fish were pit-tagged when and fin-clipped for later genotyping. Genotyping and genotypic sex of the parent and offspring were performed by genotyping-by-sequencing using 177 nucleotide polymorphisms (SNPs) which included *vgll3* and *six6* SNPs, which were also used for parentage assignment (see, Debes *et al,* 2021, for details). The rearing temperature of the fish in this study followed a seasonal cycle (range: 4.1– 16.0°C) and the natural light regime of the latitude (61.054° N) as described in Debes *et al* (2019). In August 2018, the fish were individually PIT-tagged and a fin clip sample was taken to identify the families via genetic pedigree reconstruction. Fish were fed ad libitum except for a short period of restricted feeding for half of the tanks, which included ad libitum feeding for two days and then no feeding for five days, each week for a five-week period (Debes *et al*, 2021).

### Landmarking and trait measurements

Morphological data were collected as part of a larger effort in December 2018. Fish of both sexes were euthanized using an overdose of tricaine methanesulphonate bath, and stored briefly on ice until imaging was performed the same day. Each fish was sliced ventrally to collect data on gonad development for another study. The fish was placed on graphical paper, and the body image was taken using a digital camera (Canon PowerShot S110) mounted on a tripod (Hama Star 61), with aperture-priority exposure at F:8, with autofocus option (Supplementary Figure 1). Cross sections were obtained by slicing the fish at the posterior end of the dorsal fin (hereafter termed as large cross section) and adipose fins (hereafter termed as small cross section), respectively. These were placed on a transparent sheet and digitized using a scanner (Epson XP-305) with a modified lid to accommodate the depth of the cross sections (Supplementary Figure 1).

Landmarks were placed on the whole body and the two cross sections in a randomized order using imglab software (King, 2009), by two of the co-authors (LG for landmarked body planes, and SC for landmarked cross sections) who were blind to any information related to the genotypes or sex of the fish (Supplementary Figure 2). Landmarks were not placed around the gut region due to the previous ventral slicing. Some landmarks were solely used in the Procrustes analysis to quantify shape, while others were used to quantify specific trait values by calculating the distance between landmarks (Supplementary Figure 2, Table 1). Traits that were measured twice along the transverse plane (both the left and right sides, Table 1) were averaged prior to analysis. All measurements were reported in centimeters (cm).

**Table 1:**
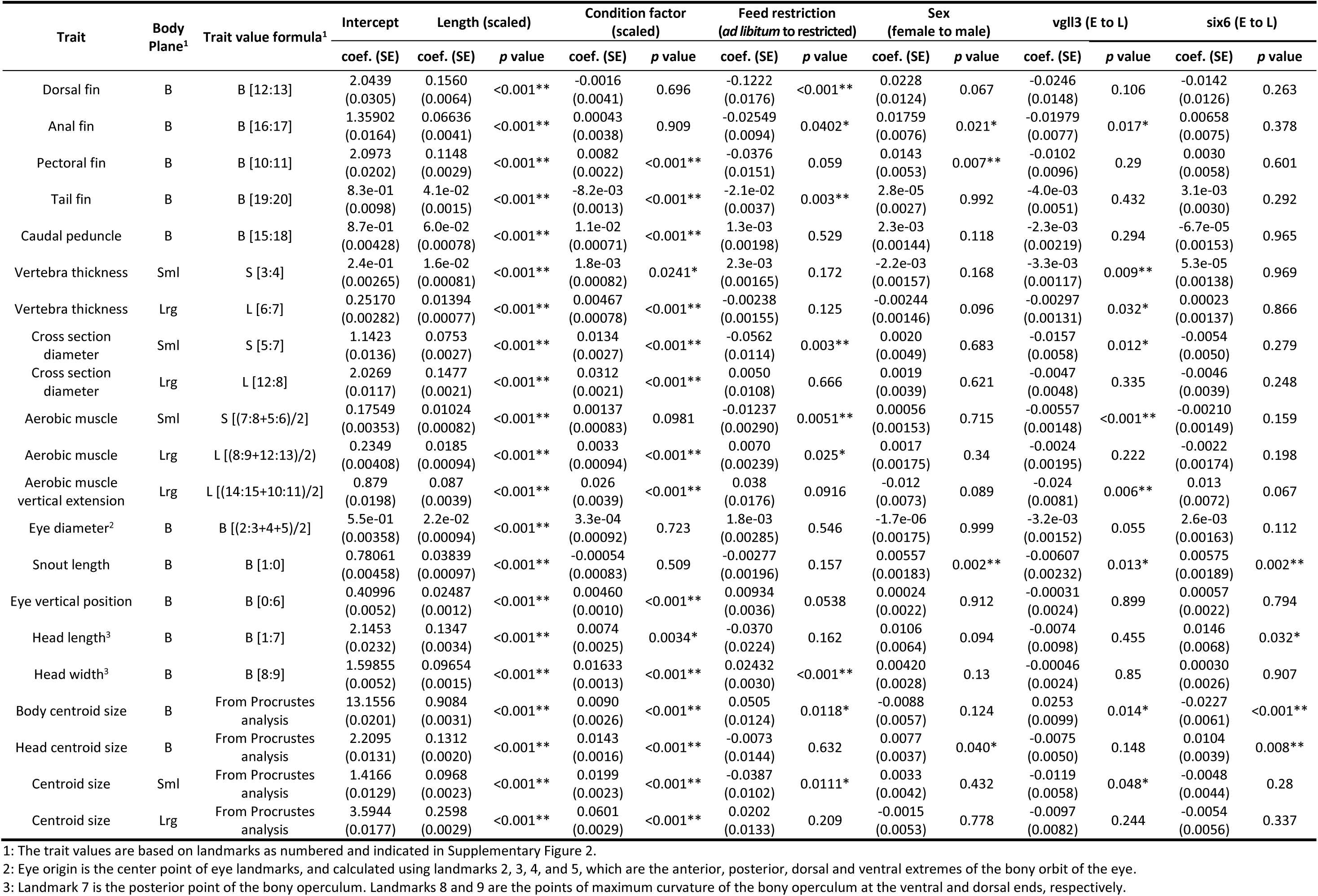
Estimated fixed effect coefficients for 21 morphological traits obtained across three body planes. Trait lengths are given in centimeters (cm) for all traits. Coef. (SE) indicates coefficients and their standard error. B, S, and L in the body plane column indicates lateral view of the body, small and large cross sections, respectively. Significant *p* values are indicated with asterisk (* and ** denotes *p*<0.05 and *p*<0.01, respectively).

### Data filtering

At the time of sampling, fish exhibited a bimodal length distribution which is often observed within juvenile salmon from the same cohort (Hoar, 1988). At the time of sampling a majority of fish used in the analysis (i.e. upper mode of size distribution, see below) had a complete smolt appearance (base on the disappearance of parr marks in the skin and silvery coloration, e.g., Debes *et al*, 2020), or were at late stages of smoltification (faint parr marks with close to silvery skin coloration). We excluded a small number of fish in the lower mode to avoid potential interactions associated with this size bimodality (fish fork length < 9 cm, N=69, Supplementary Figure 3). In addition, fish with missing *vgll3* and *six6* genotypes, and sex (N=2, N=12, and N=4, respectively), and a small number of precociously matured males (N=19) were excluded from the analysis. Missing landmarks (0.02% of the total landmarks) were imputed using the *estimate.missing* function in the *geomorph* package (version 4.0.6, Baken *et al*, 2021). A landmarked image was excluded from the analysis if more than one landmark was identified as an outlier in the Procrustes analysis (i.e. > 4 sd away from the mean). These landmarks were not corrected manually, to avoid introducing among technician variation. After data filtering, 821 individuals remained in the dataset, of which 731, 737, and 772 individuals were in the small cross section, large cross section, and whole body datasets, respectively (Supplementary Table 1). Within each dataset 16, 18, and 712 individuals were landmarked more than once to estimate repeatability. Landmarking for the first 30 and 50 images per researcher for body and cross sections, respectively, were discarded as training sets.

### Trait measurements

We quantified variation in 21 traits across various body planes with potential functional significance using the landmark positions (Supplementary Figure 2) and as formulated in Table 1.

#### Rayed fins and caudal peduncle length

We measured the length of dorsal, anal, and tail fins, between interior and exterior insertion points, as well as the caudal peduncle (Table 1, Supplementary Figure 2). The variation in these traits may be associated with habitat use, feeding, and swimming performance in salmonids and other fishes (e.g., Jonsson and Jonsson, 2011; Keeley *et al*, 2005). Increased dorsal and pectoral fin sizes in fish are likely important for increased maneuverability and stability, and likely important adaptations for faster flowing streams (Webb, 1984). Specifically, large paired fins (such as pectoral and pelvic fins) in Atlantic salmon are associated with populations that occupy faster flowing and colder streams than slower flowing and warmer rivers, an adaptation attributed to providing effective stability under fast flowing, and turbulent currents (Riddell and Leggett, 1981; Webb, 1988). In Coho salmon, juveniles of interior populations with longer migration distances appear to have smaller fins (Taylor and McPhail, 1985), which may help to reduce drag in long migrations (Varian and Nichols, 2010).

#### Eye diameter and position, and snout length

The size of the eye was approximated from the diameter, and the snout length was measured as the distance between the center of the eye and the anterior extreme of the head (Table 1, Supplementary Figure 2). These traits are morphological features associated with trophic specialization and habitat use across fish taxa (Elmer *et al*, 2010; Peris Tamayo *et al*, 2020; Snorrason, 1994).

#### Aerobic muscle

We approximate the area of aerobic muscle fibers (also called slow-twitch muscle, or red muscle) by measuring the landmark distances in the cross sections from the lateral line towards the transverse sections, and by measuring the extent of the red muscle along the lateral line (Table 1, Supplementary Figure 2). Aerobic muscle provides the energy required during steady, cruise swimming, and helps with thermal acclimation in fishes (Altringham and Ellerby, 1999; Shuman and Coughlin, 2018). Recently, it has been shown that both *vgll3* and *six6* loci are associated with whole body aerobic activity (Prokkola *et al*, 2022), which is a critical physiological process in the salmonid life cycle (Eliason and Farrell, 2015), which may indicate that aerobic muscle mass, as a proxy for aerobic activity, may be associated with these loci.

#### Cross sectional centroid size and diameter, and vertebral thickness

We measured cross sectional centroid size and horizontal diameter from both large and small cross section images (Table 1, Supplementary Figure 2), as an approximation for fusiform body shape. A slimmer and more torpedo-shaped body provides improved energy efficiency during cruising and in high flow streams by reducing water drag, and it is an important life history adaptation among fishes (Crossin *et al*, 2004; Jonsson and Jonsson, 2011a; Taylor and McPhail, 1985; Webb, 1988). In contrast, a robust cross section facilitates burst swimming, which is advantageous for prey avoidance and food acquisition (Taylor and McPhail, 1985).

We also quantified vertebral width as a proxy for cone diameter, to approximate vertebra’s flexibility/stiffness during the lateral bending movement in the undulated motion, which helps to maintain steady swimming at the expense of maneuvering capabilities (Baxter *et al*, 2022; Donatelli *et al*, 2021; Long and Nipper, 1996).

#### Head and body size

We quantified head length and width (Table 1, Supplementary Figure 2), which are important determinants of ecotypic divergence via resource and habitat use in fishes (Peris Tamayo *et al*, 2020; Simonsen *et al*, 2017). A larger head and larger snout length may be associated with a feeding regime towards piscivory (Keeley *et al*, 2005), or generally speaking, consuming larger prey items. We also measured head and body centroid sizes as an approximation for the area of the morphometric planes, which is the square root of the sum of squared distances of all the landmarks used in the Procrustes analysis from the point of center of gravity (Klingenberg, 2016). The Atlantic salmon has a smaller head-body ratio compared to the brown trout (Jonsson and Jonsson 2011), which likely makes salmon more efficient swimmers both in strong freshwater currents and during longer marine migrations.

### Statistical analysis

Our analysis framework comprised a series of analyses whereby Procrustes shape variables, and morphological traits were used as response variables. These analyses were conducted independently for the body, small cross section, and large cross section. In all analyses, length (centered to zero and scaled to have unit standard deviation), body condition (Fulton’s condition factor, scaled), sex, and food treatment were used as covariates. In all models, we implemented genetic variation as additive, where *vgll3* and *six6* genetic effects are coded as 0, 1, and 2 for EE, EL, and LL genotypes respectively (E and L refer to the alleles associated with early and late maturation, respectively), so that a unit change in genetic effect represents an additive allelic substitution effect from E to L. Unless otherwise noted, all statistical analyses were performed in R software version 4.4.1 (R Core Team, 2019).

We performed multivariate generalized Procrustes analysis to quantify shape variation (Gower, 1975), using the default settings in the *gpagen* function in the geomorph package (version 4.0.8, Baken *et al*, 2021). This function outputs Procrustes shape variables, centered on the mean value for each landmark and normalized for centroid size. Centroid size is the square root of the sum of squared distances of landmarks from the center of gravity (i.e., centroid) and it is highly correlated to the size of the fish.

Procrustes shape data was then used to perform a Procrustes linear regression analysis using the *procD.lm* function with default settings (i.e. residual randomization was used for significance testing and type III marginal sums of squares for cross products computations). The model parameters were as follows:

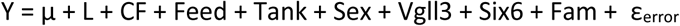

Where Y is the multivariate data frame consisting of x and y coordinates for Procrustes shape coordinates of landmarks, μ is the intercept, L is the scaled fork length, CF is the scaled Fulton’s condition factor, and Feed is the factor associated with feed treatment (ad-libitum vs restricted feed). Other terms are the rearing tank ID that the fish were reared after the swim-up stage (Tank), sex (Sex), family (Fam), and the additive genetic effects for *vgll3* and *six6*, respectively. Since the *procD.lm* function does not accommodate random terms, we included tank and family effects as fixed terms. Trait values were averaged prior to the analyses when an individual had replicate measurements. We reported multivariate analysis of variance Pillai’s trace statistics, estimated using the *manova.update* function in the RRPP package (version 2.0.3, Collyer *et al*, 2018), and plot individual landmark differences between loci using the *plotRefToTarget* in the geomorph package.

To decompose the variation associated with trait values and estimate parameter coefficients, we employed a linear mixed effect (animal) model using the *relmatLmer* function in lme4qtl (version 0.2.2, Ziyatdinov *et al*, 2018), which is an implementation of the *lmer* function in the lme4 package (Bates, 2010), and can fit the animal model (i.e., accommodate the additive relatedness matrix). When estimating fixed coefficients, the model was fitted with maximum likelihood using the *REML* = *FALSE* option in the *relmatLmer* function. The model structure was as follows:

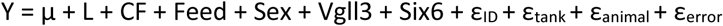

Fixed effect terms are as described above except that the fixed tank and family terms in the previous model were replaced with random tank and animal terms, respectively. Random terms thus included, ε_ID_ which are the individual effects, ε_tank_ are the effects associated with the tank effects, and ε_error_ is the measurement error. ε_animal_ is the random term associated with the background genetic effects after controlling for *vgll3* and *six6* fixed effects. The relatedness matrix was obtained using the pedigree data in Debes *et al* (2019) using the *getA* function in the pedigreem package (version 0.3.4, Vazquez *et al*, 2010). Since we used repeated measurements in the analysis, ε_ID_ reflects the error term, and ε_error_ is the residuals that quantifies deviations between repeated samples (i.e. measurement error).

The significance of the fixed effects terms was estimated using the *as_lmerModLmerTest* function in the lmerTest package (version 1.1-3, Kuznetsova *et al*, 2017) in R using type 3 F test with Satterthwaite’s method to estimate the effective degrees of freedom.

We then estimated the additive genetic variation, the contribution of *vgll3* and *six6* loci to the additive genetic variation, evolvability, and repeatability. For these, we employed three models similar to above, but the first model (Model_VA.full_) did not contain *vgll3* or *six6* to estimate total additive genetic variation, and was fitted using the *REML* = *TRUE* option in the *relmatLmer* function to obtain unbiased estimates of variance components. As such, the variance of ε_animal_ (*V*_animal_) in the Model_VA.full_ is the total additive genetic variation (*V*_A_). Percent contribution of *vgll3* and *six6* are calculated as%*V*_A.vgll3_ = 100 * ((2 x L_vgll3_ x E*_vgll3_* x (β*_vgll3_*^2^ – se(β)*_vgll3_*^2^) / *V*_animal_), and %*V*_A.six6_ = 100 * ((2 x L *_six6_* x E*_six6_* x (β *_six_6*^2^ – se(β) *_six6_*^2^))) / *V*_animal_), as in (Luo *et al*, 2003), where L*_vgll3_*, E*_vgll3_*, L *_six6_* and E*_six6_* denotes frequency of L and E alleles in *vgll3* and *six6* loci, respectively, β*_vgll3_* and β*_six6_* denotes *six6* and *vgll3* E to L substitution effects, respectively, and se denotes standard error of the estimate. Percent repeatability for each model is calculated as R = *V*_ID_ / (*V*_ID_ + *V*_error_) * 100, and heritability values were calculated as *h*^2^ = *V_animal_* / (*V*_animal_ + *V*_tank_ + *V*_ID_), where *V*_tank_, *V*_ID_, and *V*_error_ are the model variance estimates associated with tank (ε_tank_), individual (ε_ID_), and error (ε_error_) terms in Model_VA.full_. Similarly, evolvability (*I*_A_) was calculated as *V*_A_ / ̅*y*^2^ using Model_VA.full_, where ̅*y* is the mean trait value (Hansen *et al*, 2011; Houle, 1992). Evolvabilities are interpreted as expected proportional responses in the trait mean under unit selection (i.e. the selection on fitness itself; Hansen *et al*, 2003; Hansen *et al*, 2011). For all variance components, 95% confidence intervals were estimated by obtaining the posterior distributions of random effect terms in the model (Model_VA.full_), using the *sims* function in the arm package(version 1.1.4-4, Gelman and Su, 2020) in R, using 10000 permutations. The simulations generate 10000 estimates for random effect terms, which are then used to calculate the variance components as above, with 250^th^ and 9750^th^ ranked estimates recorded as 95% CIs. We did not control for the effect of dominance in the analyses, which may have inflated the estimates of additive genetic variance (Lynch and Walsh, 1998).

## Results

### Shape analysis

The multivariate analyses using Procrustes shape variation as the response showed that genetic variation in *six6*, but not in *vgll3*, is associated with shape change in the body plane (Table 2). The change in the body plane appears to be related to changes in the head region associated with *six6*, whereby a substitution from *six6*^E^ to *six6*^L^ elongates the head and slightly bends up the lateral body positioning at ends of the lateral line. In contrast, *vgll3*^E^ to *vgll3*^L^ substitution reduces the head area relative to the body area and pushes the anterior anal fin landmark toward the posterior end of the body plane, albeit the effect is statistically insignificant (Figure 1a). Genetic variation in *vgll3* and *six6* are not associated with overall shape changes in small and large cross sections. Although not statistically significant, a *vgll3*^E^ to *vgll3*^L^ substitution was linked with a narrower horizontal width, a smaller aerobic muscle area in the small cross section (Figure 1b), and a narrower aerobic muscle extension perpendicular to the lateral line in the large cross section (Figure 1c). These results are concordant with the trait-based analysis (see below). Overall, family, tank, feed restriction, and length were important determinants of shape variation in all planes, and condition factor was also significant in the analysis of the body plane (Table 2). Sex was an insignificant determinant of shape in all three body planes (Table 2).

**Figure 1:**
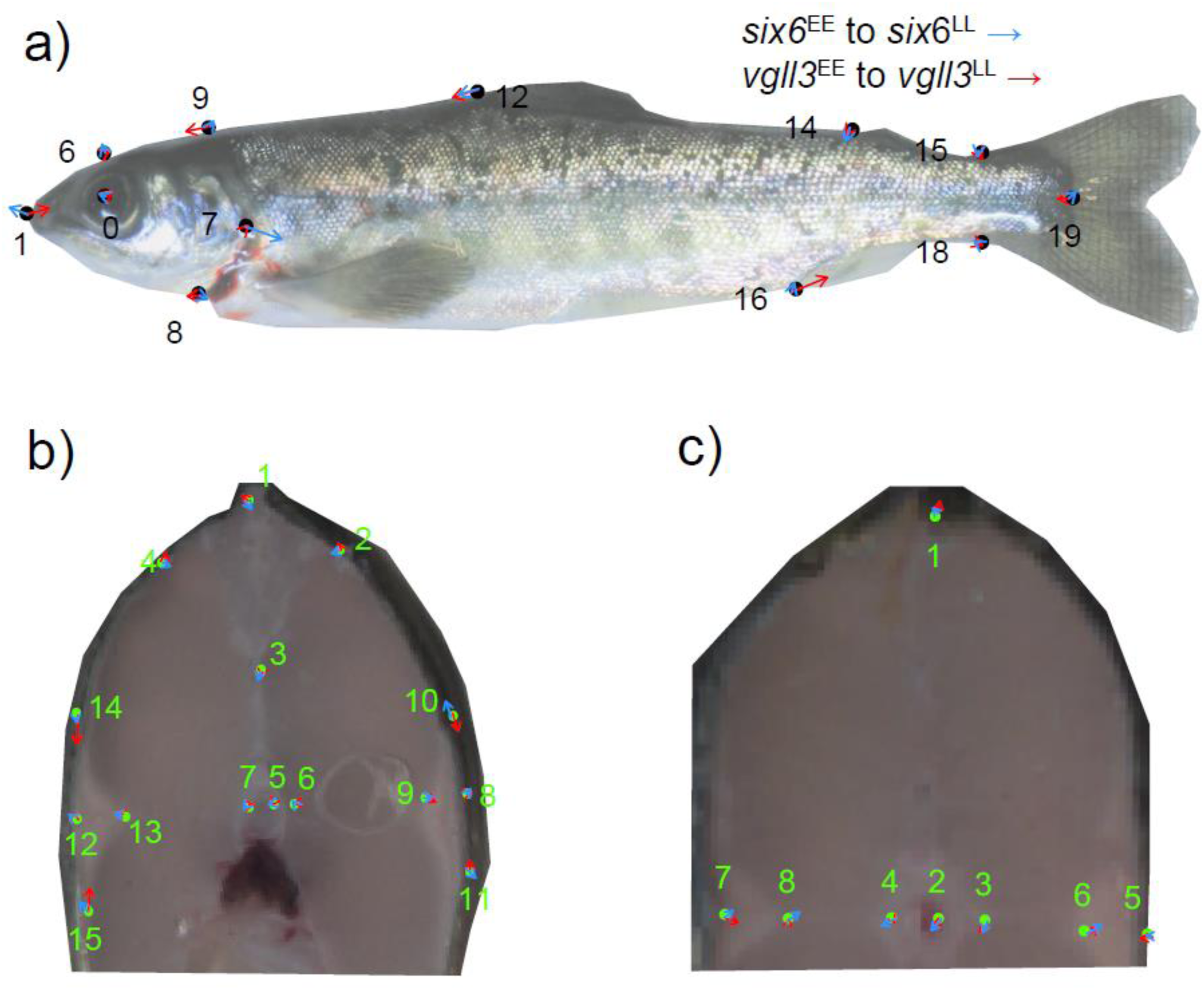
The difference between *vgll3*^EE^ and *vgll3*^LL^ (blue lines), and *six6*^EE^ and *six6*^LL^ (red lines) genotypes on body shape of juvenile Atlantic salmon using thin-plate spline in the body (a), and large (b) and small (b) cross sections. The black dots in a, and green dots in b and c are the predicted shape coordinate associated with *vgll3*^EE^ and *six6*^EE^ after the generalized Procrustes analysis. For all images, the effect sizes are magnified by 10x. Landmark numbers are based on Supplementary Figure 2. Note that the body and cross section images are to illustrate the location of landmarks on these body planes, and were slightly distorted to fit onto the average coordinates obtained by Procrustes analysis.

**Table 2:**
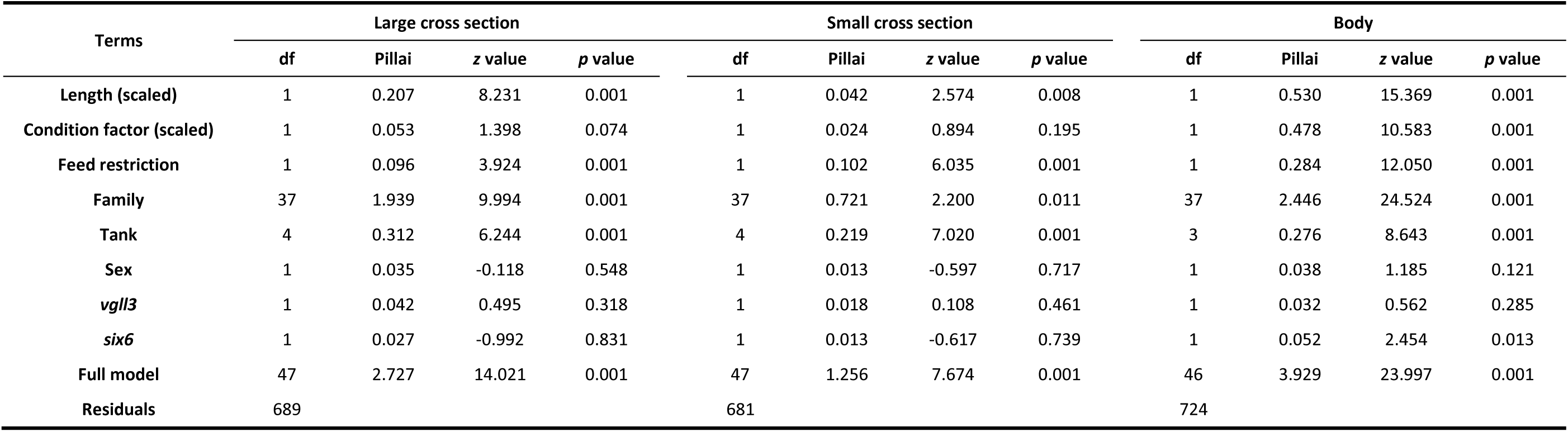
Multivariate analysis using Procrustes shape variable across three body planes in juvenile Atlantic salmon. Pillai is the Pillai’s trace test statistic to evaluate each term’s significance in the multivariate analysis of variance.

#### Trait analysis

After correcting for body length, condition factor, feed restriction, and sex, nine out of 21 morphological traits were significantly associated with the *vgll3* locus, while four were associated with *six6* locus (Table 1, Figure 2. See also Supplementary Table 3 for *t*-values and effective degrees of freedoms of terms). In general, the *vgll3*^E^ to *vgll3*^L^ substitution effect decreased trait values (Table 1, Figure 2) with changes in trait values between *vgll3*^EE^ and *vgll3*^LL^ genotypes being > 1%, and up to 3% of the mean trait values (Figure 2). Notably, the aerobic muscle depth in the small cross section was reduced by 3.45% (*p*< 0.001) and the aerobic muscle vertical extension in the large cross section was reduced by 2.72% (*p* = 0.006) by the *vgll3*^E^ to *vgll3*^L^ allelic substitution (Table 1, Figure 2). The *vgll3* allelic substitution effect associated with lateral red muscle length was statistically significant when three aerobic muscles trait values were combined into one composite value (*p*< 0.001, Supplementary Table 4). The *vgll3*^L^ allele also decreased vertebral thickness in both small (1.37% decrease, *p* = 0.009) and large (1.21% decrease, *p* = 0.032) cross sections (Table 1, Figure 2). The *vgll3*^E^ to *vgll3*^L^ substitution effect significantly reduced the small cross section diameter by 1.43% (*p* = 0.012), whereas the small section centroid size was reduced by 0.86% (*p* = 0.012, Table 1, Figure 2). Reduced eye diameter, snout length, and anal fin length were also associated with the *vgll3*^L^ allele (Table 1, Figure 2). The trait values for all ray fins (dorsal, anal, pectoral, and tail fins) was trending toward a smaller size in relation to the late maturing genotype (Table 1, Figure 2), and their cumulative effect, i.e., effect size of all trait values combined, was statistically significant (*p* = 0.022, Supplementary Table 4). We observed that in some traits the variation explained by *vgll3*^E^ to *vgll3*^L^ substitution was up to 50% of the variation explained by length (a factor with strong, positive allometric effect on other trait values, see Table 1), suggesting that this variation is biologically important (Supplementary Figure 4).

**Figure 2:**
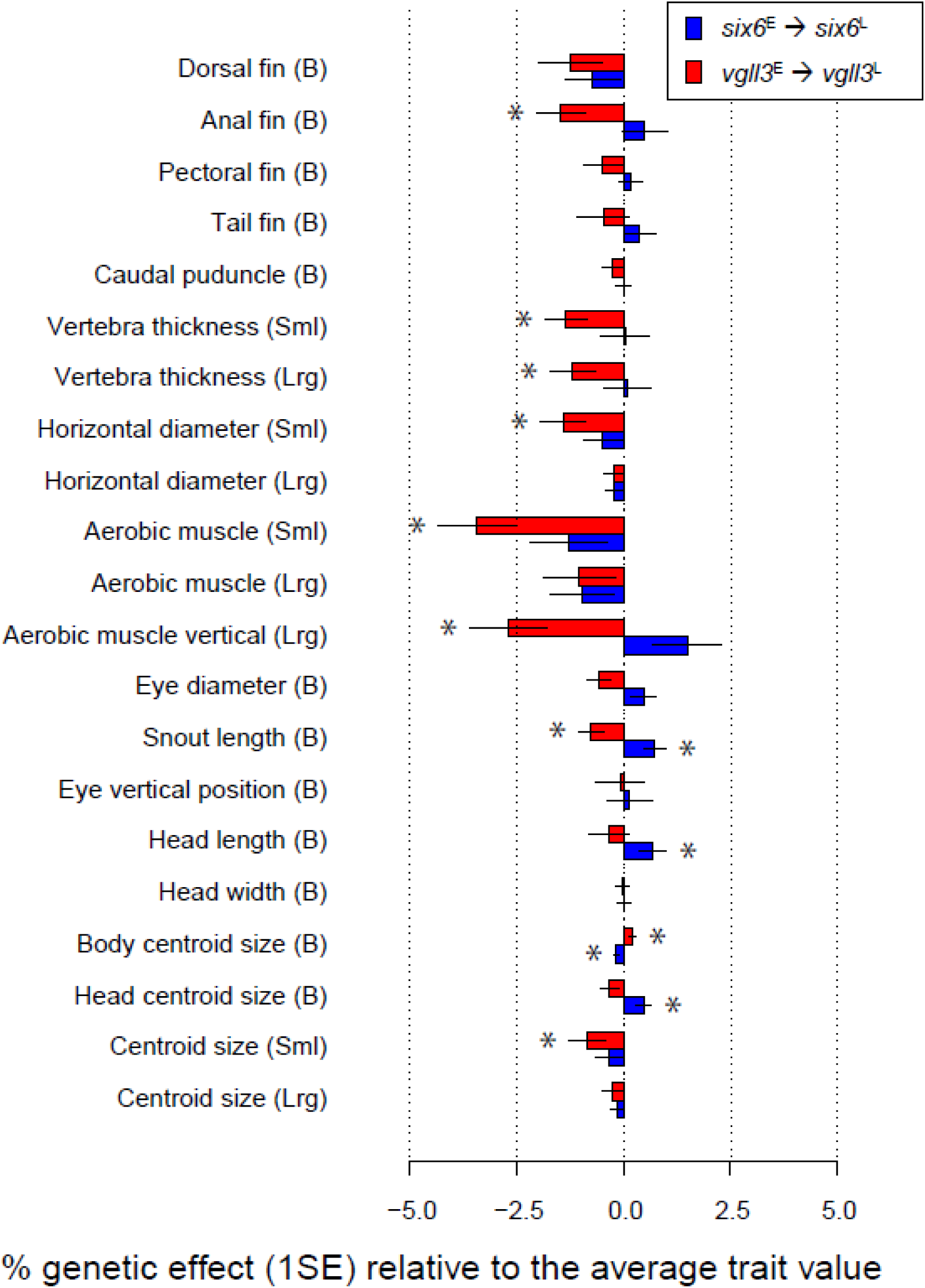
Percent trait value change associated with *vgll3*^E^ to *vgll3*^L^ (red lines), and *six6*^E^ to *six6*^L^ (blue lines) allelic substitutions relative to the average trait values. Asterisks denote significant genetic effect. Error bars and 1SE in the x-axis indicate one standard error.

Most differences explained by variation in the *six6* genomic region were related to changes in head and body size. The *six6*^E^ to *six6*^L^ substitution was associated with a 0.17% decrease in body centroid (*t* = –3.70, *p*< 0.001) and a 0.47% increase in head centroid size (*t* = 2.68, *p* = 0.008, Table 1, Figure 2). The snout length, measured as the distance between the center of the eye and the maximum anterior point in the body, was elongated by 0.73% in fish with an additional *six6*^L^ allele (*t* = 3.05, *p* = 0.002, Table 1, Figure 2). Head length appears to be a significant determinant of changes in head structure, the effect of an additional *six6*^L^ allele explained a 0.33% increase in average trait value (*t* = 2.15, *p* = 0.032, Table 1, Figure 2).

Sex effects typically ranged from smaller and up to the same order of magnitude as the allele substitution effects in the life-history loci (Table 1). Notably, the snout, pectoral fin, and anal fin were significantly larger in males compared to females (*p* = 0.002, *p* = 0.007, and *p* = 0.021, respectively. Table 1). In addition, the head centroid was larger in males compared to females (*p* = 0.040, Table 1). As expected, length was a strong predictor of all traits, and condition factor was also positively associated with most traits (Table 1). Feed restriction was notably associated with a decrease in fin trait values, such as the dorsal fin (*p* < 0.001), anal fin (*p* = 0.040), and tail fin (*p* = 0.003, Table 1). Interestingly, feed restriction substantially reduced the centroid size (*p* = 0.011), cross section diameter (*p* = 0.003), and the size of the aerobic muscle (*p* = 0.005) in the small cross section, but not in the large cross section (Table 1).

As length is a strong predictor of both trait values and overall body shape, a spurious association between traits and the life-history loci may be expected if these loci affect trait values via length as a mediator. To test this, we constructed a model to include the interaction between length and life-history loci. Across 21 traits, we did not observe a meaningful difference in main genetic effects, nor were interaction term *t*-values substantial (Supplementary Figure 5), suggesting that length-by-genotype interaction did not drive the estimated relations.

*Heritability, evolvability, and additive genetic* variation: Trait heritabilities were low to moderate in most cases, with a mean heritability of 0.19 across all 21 traits (Table 3). Length of tail fin and body centroid size exhibited the highest heritability with 0.50 (95% CI 0.47-0.54) and 0.47 (95% CI 0.43-0.50), followed by the pectoral fin with 0.41 (95% CI 0.36-0.45). The repeatability of trait measurements was moderate (mean repeatability 73.0%, Table 3). The relatively low repeatability than expected (Takacs *et al*, 2016) is elevated likely due to low intra-population variation obtained in the common garden setting relative to most morphometrics studies that focuses on larger biological variation, e.g. between ecomorphs of conspecifics or across species, but complex trait measurements combined with suboptimal fish imaging using gutted individuals, and measurer’s expertise may also contribute to such variation (Vrdoljak *et al,* 2020, Takacs *et al*, 2016).

**Table 3:**
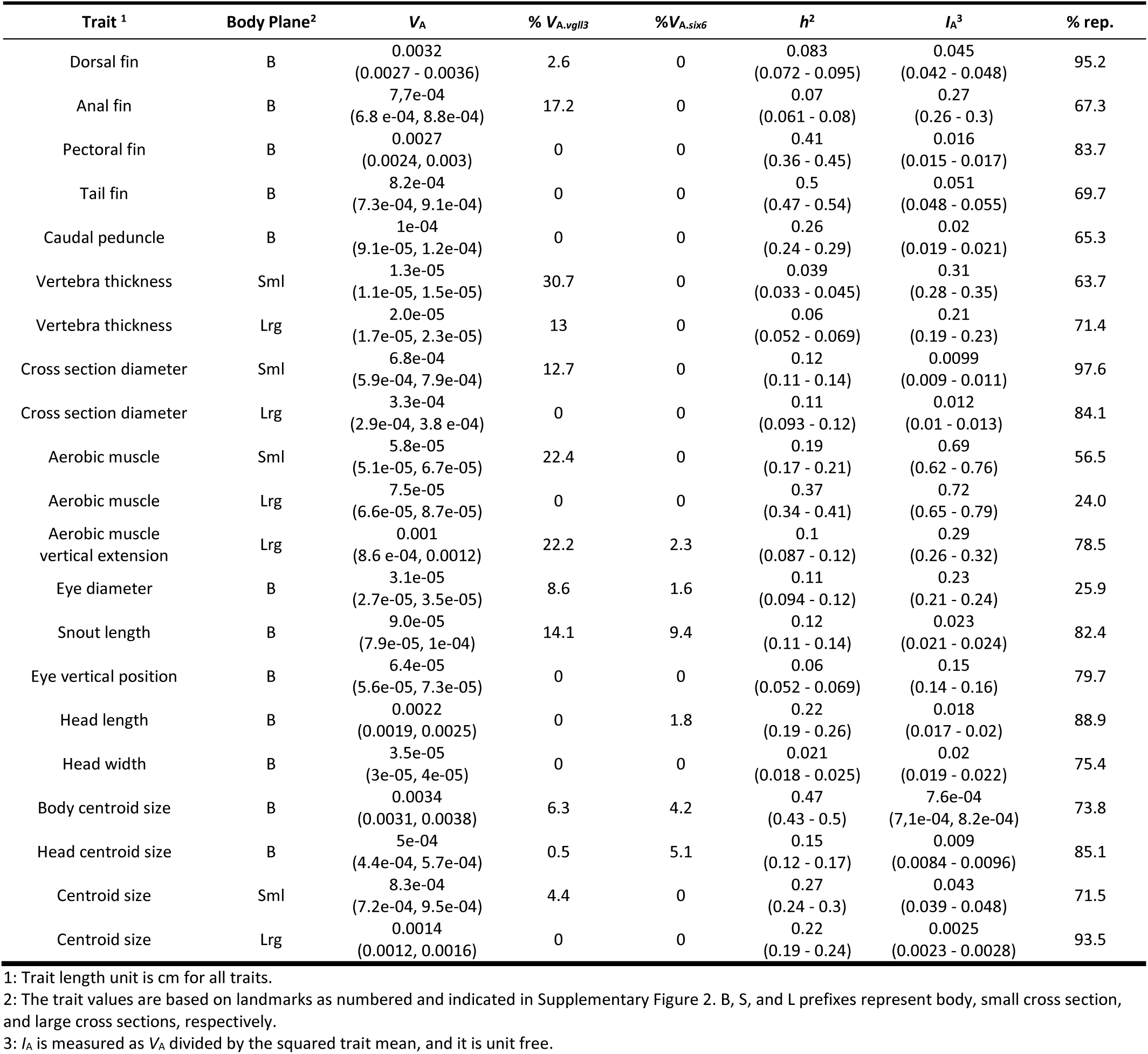
Variance components for 21 morphological traits obtained from three body planes. *V*_A_, *h*^2^, and *I*_A_ are additive genetic variance, heritability, and evolvability, respectively, after controlling for length, condition factor, feed restriction, and sex.% *V*_A.vgll3_ and%*V*_A.six6_ are additive genetic variation explained by the *vgll3* and *six6* loci, respectively (values are truncated to zero if the estimate is less than zero). Values indicate medians, with 95% CI given in parenthesis. % rep. indicate the percent repeatability estimated from repeated measurements of individuals, and to assess the contribution of measurement error to trait measurements.

Evolvabilities (mean-scaled additive genetic variances, denoted as *I*_A_) differed greatly among traits (Table 3). Interestingly, body centroid size had the lowest evolvability (<0.001), meaning that even under unit directional selection (the strength of selection on fitness itself, Hansen *et al,* 2003), the expected increase in this trait value would be less than 0.1%. On the other hand, large cross section aerobic muscle had an evolvability as high as 0.72, meaning that the expected increase in this trait under strong selection could be as high as 71% of the mean trait value. In general, out of 21 traits measured, aerobic muscle and vertebrae thickness traits had higher *I*_A_, while centroid sizes and cross section diameters had lower *I*_A_ (Table 3).

Additive genetic variation explained by the *vgll3* locus was non-zero for 12 traits (Table 3). Consistent with the fixed effect parameterization of the genotype, a substantial portion of the additive genetic variation in vertebra thickness was explained by *vgll3*, with 30.7% and 13.0% for small and large cross sections, respectively (Table 3). A significant portion of the additive genetic variation in small section aerobic muscle, cross section diameter, and body centroid was also explained by the *vgll3* locus, with 22.4%, 12.7, and 6.3% for each trait, respectively (Table 3). The contribution of *six6* locus to additive genetic variation associated included snout length (9.4%), and head and body centroid size (5.2% and 4.2%, respectively, Table 3) consistent with the locus effect size.

## Discussion

Intraspecific differences in functionally important morphological traits may provide important insights into the mechanistic basis of adaptation. We detected several morphological traits in juvenile Atlantic salmon that were linked with variation in genomic regions previously reported to be associated with life-history diversity and known to be under local selection. The genetic polymorphism in the *vgll3* locus appears to shape a suite of traits associated with swimming performance, while the genetic polymorphism in the *six6* locus shapes head and body size, and head features such as snout length, which potentially affects resource acquisition efficiencies for different foraging strategies. On the other hand, these genomic regions have only a small effect on the general shape of the morphological planes. The only statistically significant effect was detected in the body plane plan that was associated with the *six6* loci (Table 2), for which the most striking shape change at the vector level elongates the head length (in relation to *six6*^E^ to *six6*^L^ substitution, Figure 1), which is concordant with change in the head/body proportions observed in the univariate trait analysis. Our results support these genomic regions having broad phenotypic effects, which may have emerged from either a form of pleiotropy or linkage disequilibrium. Our design does not allow us to categorize the pleiotropic effects further, whether it is based on an effect of single causal variant (biological pleiotropy), or based on effects associated with different phenotypes along a biochemical pathway (mediated pleiotropy), or as a result of linkage disequilibrium between two independent causative loci (spurious pleiotropy, e.g., Solovieff *et al*, 2013). Yet, the strong genetic interdependence among traits associated with *six6* and *vgll3* may have a developmental origin, which is supported by the patterns of gene expression in these genomic regions during development (Moustakas-Verho *et al*, 2020; Kurko *et al*, 2020), and their molecular regulatory functions, (Ahi *et al*, 2022; Kurko *et al*, 2020; Mobley *et al*, 2021;; Verta *et al*, 2020; Verta *et al*, 2024). For example, being a co-transcription factor binding to the TEAD protein, *vgll3* is likely an important regulator of the hippo pathway (Kurko *et al*, 2020) which is involves in organ growth via controlling cell growth (Huang *et al,* 2005). Likewise, allele specific *vgll3* regulation of pubertal development, interact with signaling pathways with diverse molecular functions, such as TGF-β signaling, and wnt pathway which have developmental functions during body axes formation, as well as cell fate commitment (Xue *et al,* 2025; Wu and Hill, 2009).

The suite of morphological traits associated with *vgll3* variation reflects changes along a functional axis that facilitates energy-efficient cruising (*vgll3*^L^) versus maneuvering and acceleration (*vgll3*^E^). It appears that that individuals with *vgll3*^E^ genotype have a robust build to facilitate maneuvering and acceleration, and larger aerobic activity (i.e. increased red muscle area, see also below), while individuals with *vgll3*^L^ allele have a slender body build for energy efficiency swimming (i.e. smaller cross section diameter, and smaller ray fins, see also below). Energy efficient cruising is an important trait especially in the marine environment, while better maneuvering and acceleration help coping with biotic (i.e. competition and predation risk) and abiotic (flow regime) factors in the freshwater habitat. The larger aerobic muscle area associated with the *vgll3*^E^ allele could indicate a higher potential for aerobic muscle activity. This result is concordant with (Prokkola *et al*, 2022) who demonstrated that juvenile Atlantic salmon with the early maturing genotype have a higher maximum metabolic rate (MMR) and aerobic scope (AS). Aerobic locomotion is expected to be central for Atlantic salmon during the marine phase, where salmon sub-adults migrate and continuously forage for food. Our results suggest salmon with the early genotype are more likely to maintain sustained swimming and foraging activities, beneficial during marine exploration, which may help to facilitate earlier maturation, with poor performers having to delay reproduction until they can utilize sufficient energy for return migration. On the other hand, having a slender body and energy-efficient cruising may be more advantageous in later years, since individuals with more *vgll3*^L^ alleles appears to perfrom better than indiviauls with *vgll3*^E^ allale at older sea-age groups (i.e. Barson *et al*, 2013). However, it is unclear if the genetic association between life-history loci and morphological traits persists between life stages, or across different environments. The parr-smolt transformation (smoltification) includes morphological changes that prepare juvenile Atlantic salmon for the marine environment (Jonsson and Jonsson, 2011), hence phenotype-genotype association might differ either as a result of smoltification or change in the environment (Aguirre *et al*, 2014). However, since individuals used in this study have a complete or nearly complete smolt phenotype, conclusions may be more relevant to the marine than to the freshwater life stage.

Increased vertebral thickness associated with the *vgll3*^E^ allele could provide stiffness for energy efficient undulation movement during steady swimming (Donatelli *et al*, 2021), though it may come at the expense of body flexibility that may facilitate piscivorous prey capture. A larger aerobic muscle area and thicker vertebra may eventually facilitate earlier maturation via more energy efficient resource acquisition at sea, suggesting that local adaptation for optimal age structure could be partly mediated by conditions at sea. Hence, while our results do not explain the causation of early male maturation in freshwater, we cannot explicitly rule out that the link between *vgll3* and age at maturity is mediated via its effect on body shape. Sex was also an important factor shaping the trait variation. Intriguingly, significant changes in shape associated with sex and *vgll3* genotype was also in concordant with respect to maturation timing. As such, males, which on average mature earlier than females, and the *vgll3* allele associated with early maturation had similar direction of change in these morphological traits.

Larger fin sizes in fish increase maneuverability and stability (Webb, 1984), and correspondingly in Atlantic salmon, larger fins occur in populations that occupy faster flowing streams (Riddell and Leggett, 1981). Overall, we found that the ray-fins of fish were consistently larger for the *vgll3*^E^ allele than the *vgll3*^L^ allele (Supplementary Table 4). However, this improvement for maneuverability and stability may come at the expense of efficiency in longer marine migrations, as a result of increased drag. For example, in coho salmon, juveniles of inland populations with longer migration distances have smaller fins (Taylor and McPhail, 1985), which may help reduce drag in long migrations (Dynes *et al*, 2005; Varian and Nichols, 2010). A larger cross section diameter and larger fin lengths associated with the *vgll3*^E^ allele likely facilitate maneuvering at the expense of efficient cruising, which is contrast with the *vgll3*^E^ effect that increases aerobic muscle area and vertebral thickness. Similarly, the slender body (a smaller cross section diameter) associated with *vgll3*^L^ is consistent with minimizing drag for a cruising body-form (Webb, 1988), advantageous for a multi-sea winter strategy that migrates longer distances in the ocean (e.g., O’Sullivan *et al*, 2022). On the other hand, the direction of effect on fin size associated with *vgll3*^E^ *to vgll3*^L^ substitution is concordant with the effect of the restricted-food regime, suggesting that the allelic substitution effect may be correlated to food limitation conditions. Thus, overall, the direction of *vgll3* genetic effects is not functionally consistent in relation to proposed ecological roles of these morphological traits.

The changes associated with the genetic variation in the *six6* locus that are affecting body and head sizes are linked to ecological processes related to resource acquisition mechanisms. A larger head (and a smaller body) associated with the *six6*^L^ allele and larger snout length may be associated with a feeding regime towards piscivory (Keeley *et al*, 2005), or generally consuming larger prey items. This interpretation corroborates the study by Aykanat *et al* (2020), in which fish with the *six6*^LL^ genotype were shown to be more efficient in resource acquisition than fish with the *six6*^EE^ genotype.

Evolvabilities (*I*_A_) ranged from normal to very high compared to a recent meta-analysis (Hansen *et al*, 2011). The median evolvability for length traits is 0.001 (or 0.1%, Hansen and Pélabon, 2021; Hansen *et al*, 2011). Normal evolvability values were found for centroid sizes and cross section diameters, while aerobic muscle and vertebrae thickness had high evolvabilities, suggesting that these traits could evolve rapidly in response to selection induced by environmental change. Interestingly, a substantial portion of the additive genetic variation associated with some traits, such as aerobic muscle area and vertebrae thickness, was explained by the *vgll3* locus, suggesting the locus’ importance in shaping the additive genetic landscape.

Taken together, *vgll3* and *six6* loci have complex associations with morphology. These do not consistently align with adaptive differences between early– and late-maturation genotypes associated with marine migration, but could be important e.g., during spawning where higher maneuverability of smaller males may be beneficial for reproductive success, or in early life stages. On the other hand, it is also plausible that these changes are genetically correlated as part of broad changes during development, and some of which have potentially maladaptive consequences. When traits are genetically correlated, the cumulative adaptive value of trait values cannot be optimal (Schluter, 1996), since breaking suboptimal trait combinations that are genetically correlated may be challenging for evolution (Blows, 2007). Finally, it is challenging to assess the biological significance of the scale of trait variation between loci. However, effect size appears to be large for some trait, i.e., as inferred by up to 50% effect size of allelic substitution effect relative to the effect of length (Supplementary Figure 4), indicating that the genetic variation produces some static allometry between individuals the same cohort, which may have functional and adaptive basis (Pelabon *et al*, 2014).

It is important to recognize that extrapolation to the marine stage of Atlantic salmon based on juvenile morphology may have limitations. It is unknown if the genotype-associated morphological variation is correlated between juvenile and adult stages. While there may be genetic correlation between freshwater and marine environments (Fleming *et al*, 1994; Morinville and Rasmussen, 2006; Taylor and McPhail, 1985), the morphological changes associated with adaptation to the marine environment, i.e., the parr-smolt transformation (smoltification), may break apart trait correlations, similar to how metamorphosis takes place in many animal taxa (e.g., Aguirre *et al*, 2014). Since the data in this study were composed of individuals that had completed the parr-smolt transformation, or are close to completion, conclusions are likely more relevant to marine stages than the freshwater stages. However, any adaptive inference associated with a specific life stage should be taken with caution until observed patterns are further tested and verified at different life stages. Finally, while it is reasonable to hypothesize that the observed *vgll3* and *six6* associated morphological variation is adaptively important, the fitness effects of this variation should be better assessed before any evolutionary conclusions can be made (Kingsolver *et al*, 1995).

While modelling non-additive architecture within a locus, such as dominance, overdominance, may be relevant, since the *vgll3* locus exhibits sex dependent dominance (Barson *et al*, 2015), in this study, genetic effects were modelled additively, to avoid conceptual and statistical complexities. However, when genetic effects were modelled non-additively (separately for *vgll3* and *six6*), parsimony was meaningfully improved (i.e., AIC for dominance gene effect model was 2 units smaller than the AIC for additive gene effect model) for four out of 21 traits for the *vgll3* gene, while none of the non-additive models performed better than additive models for the *six6* gene (Supplementary Figure 6a). In all three traits with significant non-additive genetic effect, a dominance model with *vgll3*^EL^ effect resembling *vgll3*^LL^ was supported (Supplementary Figure 6 b-d), suggesting a possible role of dominance shaping some of the trait values.

In conclusion, we have detected substantial morphological variation associated with two genomic regions, which are also major determinants of life-history variation and known to be under local selection. These loci appear to influence multiple, strongly correlated traits, suggestive of a broad regulatory role with impacts across the whole life cycle. How these genetic correlations are shaped – via single locus pleiotropy or close physical linkage of multiple genes – remains to be determined.

## Acknowledgment

We thank Artur Porto for discussion related to morphometric analyses, and Seija Tillanen, Jacqueline E. Moustakas-Verho, Nikolai Piavchenko for helps with sampling. This work was supported by the Research Council of Finland (grant nos. 328860, 353388, and 325964 to TA, and 314254, 327255 and 342851 to CRP), and the European Research Council under the European Articles Union’s Horizon 2020 and Horizon Europe research and innovation programs (grant no. 742312 and 101054307 to CRP). Views and opinions expressed are however those of the author(s) only and do not necessarily reflect those of the European Union or the European Research Council Executive Agency. Neither the European Union nor the granting authority can be held responsible for them. LG was supported by Erasmus+ programme. The authors declare no competing interests.

## Ethical approval

The experiments were approved by the Project Authorisation Board (ELLA) on behalf of the Regional Administrative Agency for Southern Finland (ESAVI) under experimental license ESAVI/2778/2018.

## Author contributions

TA, VLP, and GHB conceptualized the paper. TA analyzed the data with significant input from GHB and PVD. SJ and LG performed landmarking. AHH and AR prepared and scanned the cross sections. JE, and CRP oversaw the resources and infrastructure. PVD and CRP devised the breeding design and reared the fish. TA and PVD sampled the fish and digitized the images. TA drafted the MS with significant inputs from PVD, VLP, and GHB, and all coauthors contributed to subsequent drafts.

