## Supplementary Materials for "Large effect life-history genomic regions are associated with functional morphological traits in Atlantic salmon"

**Supplementary Table 1:** Family structure in relation to parental *vgll3* and *six6* genotypes and the number of full-sib individuals used in each of the three morphometric planes (small, large, body). Note that parental origin is inferred from genotype distribution of individuals for the *six6* genotype, therefore the parent of origin of genotypes are not known.

| Family ID | Sire | Dam | $Vgll3_{sire}$ | $Vgll3_{dam}$ | $Six6_{parents}$ | Small | Large | Body |
|---|---|---|---|---|---|---|---|---|
| 5 | 1268 | 1251 | LL | LL | LL, EL | 4 | 4 | 4 |
| 6 | 1229 | 1265 | EE | EE | EL, EL | 11 | 11 | 13 |
| 7 | 1229 | 1251 | EE | LL | LL, EL | 2 | 2 | 4 |
| 8 | 1268 | 1265 | LL | EE | EL, EL | 3 | 3 | 4 |
| 17 | 1282 | 1240 | LL | LL | LL, LL | 23 | 23 | 22 |
| 18 | 1249 | 1225 | EE | EE | LL, LL | 20 | 20 | 22 |
| 19 | 1249 | 1240 | EE | LL | LL, LL | 19 | 19 | 20 |
| 20 | 1282 | 1225 | LL | EE | LL, LL | 16 | 16 | 16 |
| 21 | 1289 | 1224 | LL | LL | LL, EL | 14 | 14 | 17 |
| 22 | 1269 | 1290 | EE | EE | LL, EL | 17 | 17 | 19 |
| 23 | 1269 | 1224 | EE | LL | LL, EL | 21 | 22 | 22 |
| 24 | 1289 | 1290 | LL | EE | LL, EL | 20 | 20 | 23 |
| 25 | 1242 | 1286 | LL | LL | LL, LL | 14 | 14 | 13 |
| 26 | 1231 | 1288 | EE | EE | LL, LL | 22 | 22 | 22 |
| 27 | 1231 | 1286 | EE | LL | LL, LL | 13 | 13 | 15 |
| 28 | 1242 | 1288 | LL | EE | LL, LL | 27 | 27 | 24 |
| 29 | 1245 | 1226 | LL | LL | LL, LL | 29 | 29 | 30 |
| 30 | 1228 | 1285 | EE | EE | EL, EL | 29 | 30 | 32 |
| 31 | 1228 | 1226 | EE | LL | LL, EL | 24 | 24 | 25 |
| 32 | 1245 | 1285 | LL | EE | LL, EL | 21 | 21 | 20 |
| 34 | 1237 | 1293 | EE | EE | LL, EL | 26 | 26 | 24 |
| 36 | 1230 | 1293 | LL | EE | LL, EL | 23 | 23 | 26 |
| 37 | 1247 | 1281_2 | LL | EL | LL, LL | 20 | 20 | 19 |
| 38 | 1276 | 1254 | EE | EE | EL, EL | 24 | 22 | 22 |
| 39 | 1276 | 1281_2 | EE | EL | LL, EL | 31 | 31 | 32 |
| 40 | 1247 | 1254 | LL | EE | LL, EL | 16 | 17 | 20 |
| 49 | 1256 | 1304 | LL | LL | LL, LL | 19 | 19 | 20 |
| 50 | 1259 | 1253 | EE | EE | LL, EL | 24 | 24 | 28 |
| 51 | 1259 | 1304 | EE | LL | LL, EL | 21 | 21 | 23 |
| 52 | 1256 | 1253 | LL | EE | LL, LL | 21 | 22 | 24 |
| 53 | 1284 | 1287 | LL | LL | EE, LL | 17 | 18 | 20 |
| 54 | 1246 | 1297 | EE | EE | LL, LL | 29 | 29 | 31 |
| 55 | 1246 | 1287 | EE | LL | EE, LL | 26 | 27 | 28 |
| 56 | 1284 | 1297 | LL | EE | LL, LL | 17 | 17 | 18 |
| 61 | 1291 | 1239 | LL | LL | LL, EL | 8 | 8 | 9 |
| 62 | 1216 | 1258 | EE | EE | EL, EL | 28 | 30 | 28 |
| 63 | 1216 | 1239 | EE | LL | LL, EL | 2 | 2 | 2 |
| 64 | 1291 | 1258 | LL | EE | EL, EL | 28 | 30 | 30 |
| Total | | | | | | 729 | 737 | 771 |

**Supplementary Table 2:** The number of individuals used in this study (N=822) from each family and tank combination.

| | Tank ID | | | | | |
|---|---|---|---|---|---|---|
| Family | T01 | T06 | T08 | T09 | T11 | T14 |
| 5 | 0 | 0 | 2 | 0 | 0 | 2 |
| 6 | 1 | 3 | 2 | 1 | 4 | 3 |
| 7 | 0 | 1 | 1 | 0 | 1 | 1 |
| 8 | 0 | 1 | 0 | 1 | 1 | 1 |
| 17 | 0 | 3 | 4 | 7 | 4 | 6 |
| 18 | 0 | 4 | 6 | 4 | 4 | 4 |
| 19 | 0 | 2 | 4 | 8 | 5 | 2 |
| 20 | 0 | 1 | 5 | 2 | 4 | 4 |
| 21 | 0 | 4 | 7 | 3 | 2 | 2 |
| 22 | 0 | 2 | 4 | 6 | 4 | 3 |
| 23 | 0 | 1 | 4 | 7 | 5 | 6 |
| 24 | 0 | 7 | 9 | 3 | 2 | 3 |
| 25 | 1 | 3 | 5 | 3 | 2 | 1 |
| 26 | 1 | 4 | 5 | 5 | 3 | 6 |
| 27 | 0 | 4 | 4 | 4 | 1 | 3 |
| 28 | 2 | 5 | 6 | 7 | 3 | 6 |
| 29 | 0 | 6 | 8 | 5 | 5 | 7 |
| 30 | 2 | 7 | 6 | 5 | 5 | 9 |
| 31 | 0 | 4 | 5 | 7 | 6 | 4 |
| 32 | 1 | 5 | 5 | 4 | 1 | 5 |
| 34 | 1 | 4 | 7 | 7 | 3 | 4 |
| 36 | 0 | 4 | 7 | 7 | 1 | 7 |
| 37 | 0 | 2 | 5 | 7 | 4 | 3 |
| 38 | 4 | 8 | 3 | 2 | 5 | 4 |
| 39 | 0 | 6 | 5 | 10 | 7 | 5 |
| 40 | 0 | 4 | 3 | 5 | 3 | 5 |
| 49 | 0 | 4 | 4 | 6 | 5 | 2 |
| 50 | 1 | 10 | 4 | 6 | 2 | 7 |
| 51 | 0 | 5 | 5 | 6 | 4 | 3 |
| 52 | 0 | 5 | 5 | 3 | 6 | 5 |
| 53 | 0 | 4 | 7 | 2 | 4 | 4 |
| 54 | 0 | 9 | 6 | 6 | 4 | 7 |
| 55 | 1 | 7 | 6 | 8 | 3 | 6 |
| 56 | 0 | 5 | 4 | 2 | 1 | 6 |
| 61 | 0 | 2 | 4 | 2 | 0 | 1 |
| 62 | 7 | 4 | 6 | 3 | 7 | 8 |
| 63 | 0 | 0 | 0 | 0 | 0 | 2 |
| 64 | 4 | 6 | 2 | 8 | 10 | 5 |

**Supplementary Table 3:** *T* values and estimated degrees of freedom for each trait and covariate for 21 morphological traits obtained across three body planes.

| Trait | Length (scaled) | | Condition factor (scaled) | | Feed restriction (*ad libitum* to restricted) | | Sex (female to male) | | *vgll3* (E to L) | | *six6* (E to L) | |
|---|---|---|---|---|---|---|---|---|---|---|---|---|
| | ***t* value** | **df** | ***t* value** | **df** | ***t* value** | **df** | ***t* value** | **df** | ***t* value** | **df** | ***t* value** | **df** |
| Dorsal fin | 24.256 | 787.4 | -0.391 | 1584.1 | -6.947 | 5.1 | 1.836 | 770.5 | -1.664 | 32.8 | -1.122 | 391.8 |
| Anal fin | 16.287 | 674.5 | 0.114 | 1058.9 | -2.723 | 5.2 | 2.309 | 765.0 | -2.557 | 25.1 | 0.883 | 234.0 |
| Pectoral fin | 40.143 | 827.0 | 3.770 | 1568.1 | -2.496 | 4.6 | 2.689 | 737.8 | -1.071 | 47.0 | 0.523 | 692.8 |
| Tail fin | 27.627 | 783.7 | -6.489 | 1377.3 | -5.787 | 4.5 | 0.010 | 750.3 | -0.792 | 49.9 | 1.055 | 714.1 |
| Caudal peduncle | 76.231 | 759.9 | 15.032 | 1196.2 | 0.676 | 5.0 | 1.566 | 742.4 | -1.067 | 33.1 | -0.044 | 576.9 |
| Vertebra thickness | 19.351 | 557.9 | 2.261 | 525.2 | 1.366 | 698.5 | -1.379 | 710.1 | -2.808 | 30.1 | 0.038 | 128.8 |
| Vertebra thickness | 18.049 | 652.4 | 5.999 | 641.2 | -1.537 | 726.2 | -1.665 | 728.5 | -2.263 | 28.2 | 0.169 | 192.7 |
| Cross section diameter | 28.230 | 702.9 | 5.028 | 712.1 | -4.930 | 6.0 | 0.409 | 718.3 | -2.687 | 26.2 | -1.085 | 353.4 |
| Cross section diameter | 70.170 | 721.3 | 14.812 | 730.5 | 0.466 | 4.0 | 0.495 | 725.5 | -0.980 | 28.6 | -1.157 | 409.5 |
| Aerobic muscle | 12.444 | 670.0 | 1.657 | 676.4 | -4.268 | 6.1 | 0.365 | 727.7 | -3.767 | 32.1 | -1.414 | 260.3 |
| Aerobic muscle | 19.627 | 677.0 | 3.540 | 691.8 | 2.909 | 6.5 | 0.955 | 684.7 | -1.249 | 27.8 | -1.290 | 316.0 |
| Aerobic muscle vertical extension | 22.160 | 703.7 | 6.700 | 713.9 | 2.130 | 4.6 | -1.700 | 726.7 | -2.970 | 28.3 | 1.830 | 332.0 |
| Eye diameter[2] | 23.703 | 561.3 | 0.354 | 713.4 | 0.645 | 5.4 | -0.001 | 752.5 | -2.085 | 14.7 | 1.604 | 104.0 |
| Snout length | 39.420 | 815.3 | -0.660 | 1412.8 | -1.420 | 764.6 | 3.040 | 760.8 | -2.610 | 34.6 | 3.050 | 447.2 |
| Eye vertical position | 21.097 | 768.4 | 4.481 | 1295.5 | 2.590 | 4.5 | 0.111 | 770.6 | -0.129 | 29.6 | 0.262 | 298.8 |
| Head length[3] | 39.641 | 834.5 | 2.933 | 1596.1 | -1.650 | 4.8 | 1.676 | 747.3 | -0.755 | 36.2 | 2.152 | 600.1 |
| Head width[3] | 66.275 | 618.9 | 12.331 | 1040.0 | 8.012 | 5.6 | 1.515 | 762.8 | -0.191 | 29.7 | 0.116 | 180.9 |
| Body centroid size | 293.640 | 836.1 | 3.490 | 1435.9 | 4.080 | 4.5 | -1.540 | 753.2 | 2.560 | 41.5 | -3.700 | 685.8 |
| Head centroid size | 65.766 | 828.4 | 8.950 | 1502.8 | -0.511 | 4.9 | 2.063 | 756.1 | -1.483 | 32.5 | 2.682 | 488.6 |
| Centroid size | 41.773 | 729.5 | 8.589 | 732.8 | -3.780 | 5.4 | 0.785 | 721.1 | -2.060 | 32.2 | -1.082 | 507.4 |
| Centroid size | 88.871 | 736.6 | 20.581 | 735.8 | 1.515 | 3.8 | -0.282 | 720.5 | -1.182 | 38.6 | -0.961 | 603.5 |

**Supplementary Table 4:** Estimated fixed effect coefficients for four composite morphological traits, obtained by scaling and averaging relevant trait values[1,2,3,4]. Coef. (SE) indicates coefficients and their standard error. Significant *p* values are indicated with asterisks (* and ** denote *p*<0.05 and *p*<0.01, respectively.)

| Trait | Intercept | Length (scaled) | | Condition factor (scaled) | | Feed restriction (*ad libitum* to restricted) | | Sex (female to male) | | *vgll3* (E to L) | | *six6* (E to L) | |
|---|---|---|---|---|---|---|---|---|---|---|---|---|---|
| | **coef. (SE)** | **coef. (SE)** | ***p* value** | **coef. (SE)** | ***p* value** | **coef. (SE)** | ***p* value** | **coef. (SE)** | ***p* value** | **coef. (SE)** | ***p* value** | **coef. (SE)** | ***p* value** |
| Ray fins all traits combined[1] | 0.1979 (0.1256) | 0.8161 (0.0184) | 44.33 <0.001** | -0.0306 (0.0150) | 0.042 | -0.4069 (0.0708) | 0.003 | 0.1044 (0.0339) | 0.002 | -0.1483 (0.0626) | 0.022* | 0.0365 (0.0369) | 0.322 |
| Aerobic muscle all traits combined[2] | 0.21630 (0.1152) | 0.71067 (0.0259) | 27.4080 <0.001** | 0.15292 (0.0260) | <0.001** | 0.00748 (0.0855) | 0.93 | -0.00716 (0.0481) | 0.88 | -0.19916 (0.0512) | <0.001** | -0.02533 (0.0479) | 0.597 |
| Vertebrate all traits combined[3] | 0.12698 (0.1029) | 0.66299 (0.0276) | 24.0553 <0.001** | 0.14673 (0.0278) | <0.001** | -0.00491 (0.0552) | 0.93 | -0.09784 (0.0521) | 0.061 | -0.13323 (0.0481) | 0.009** | 0.02222 (0.0497) | 0.66 |
| Cross section diameter all traits combined[4] | 0.2439 (0.0858) | 0.8382 (0.0155) | 54.073 <0.001** | 0.1675 (0.0155) | <0.001** | -0.2529 (0.0803) | 0.023 | 0.0136 (0.0285) | 0.633 | -0.0915 (0.0345) | 0.013* | -0.0388 (0.0292) | 0.186 |

1. Dorsal fin, anal fin, pectoral fin, and tail fin trait values were scaled and averaged for all individuals.
2. Small cross section - aerobic muscle, large cross section - aerobic muscle, and large cross section - aerobic muscle vertical trait values were scaled and averaged for all individuals.
3. Large cross section - vertebra width, and small cross section- vertebra width trait values were scaled and averaged for all individuals.
4. Large cross section - diameter, and small cross section diameter trait values were scaled and averaged for all individuals.

a)
b)
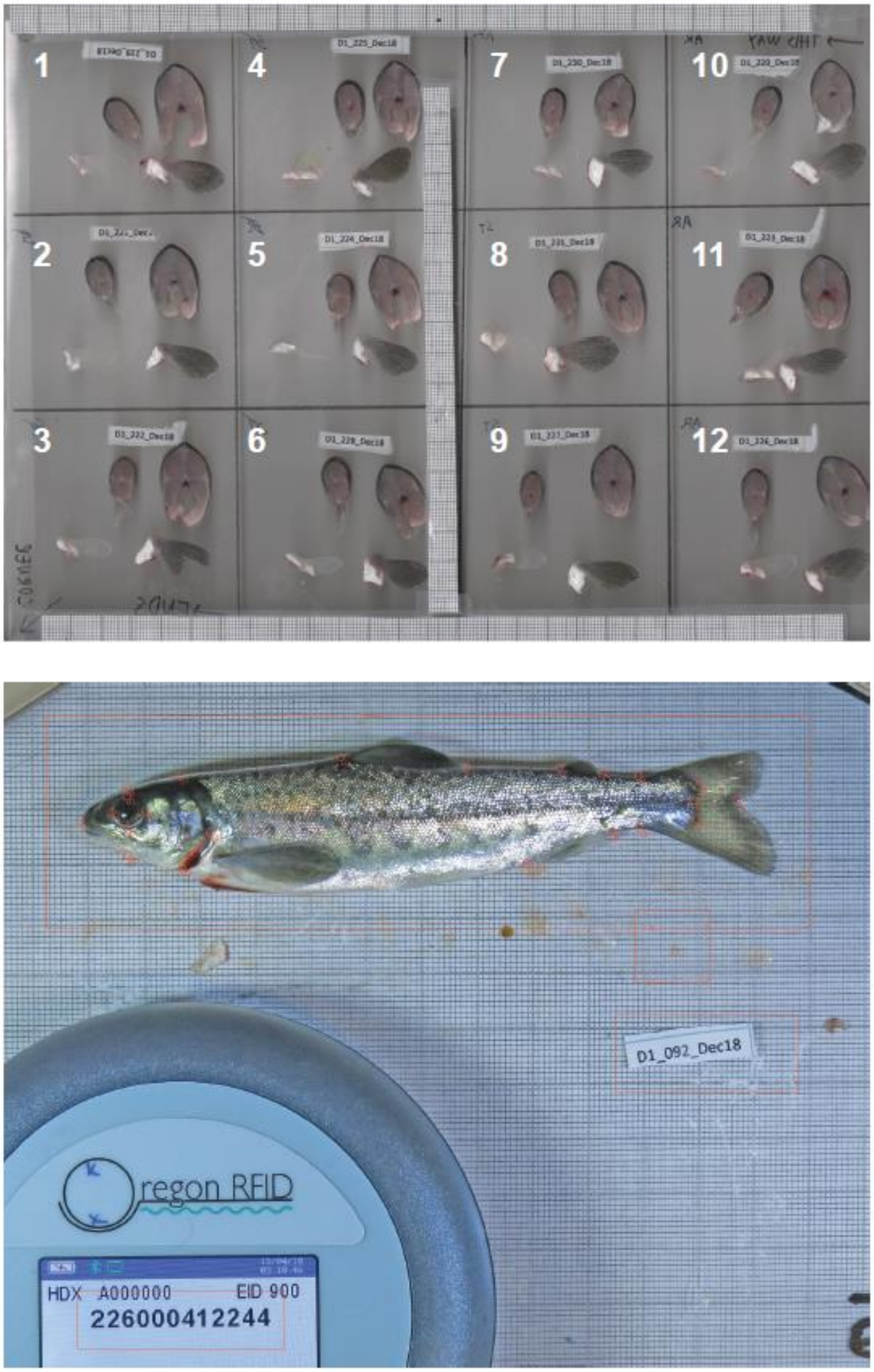


**Supplementary Figure 1**: An example of the layout of image screening of cross sections (a), and lateral view (b).

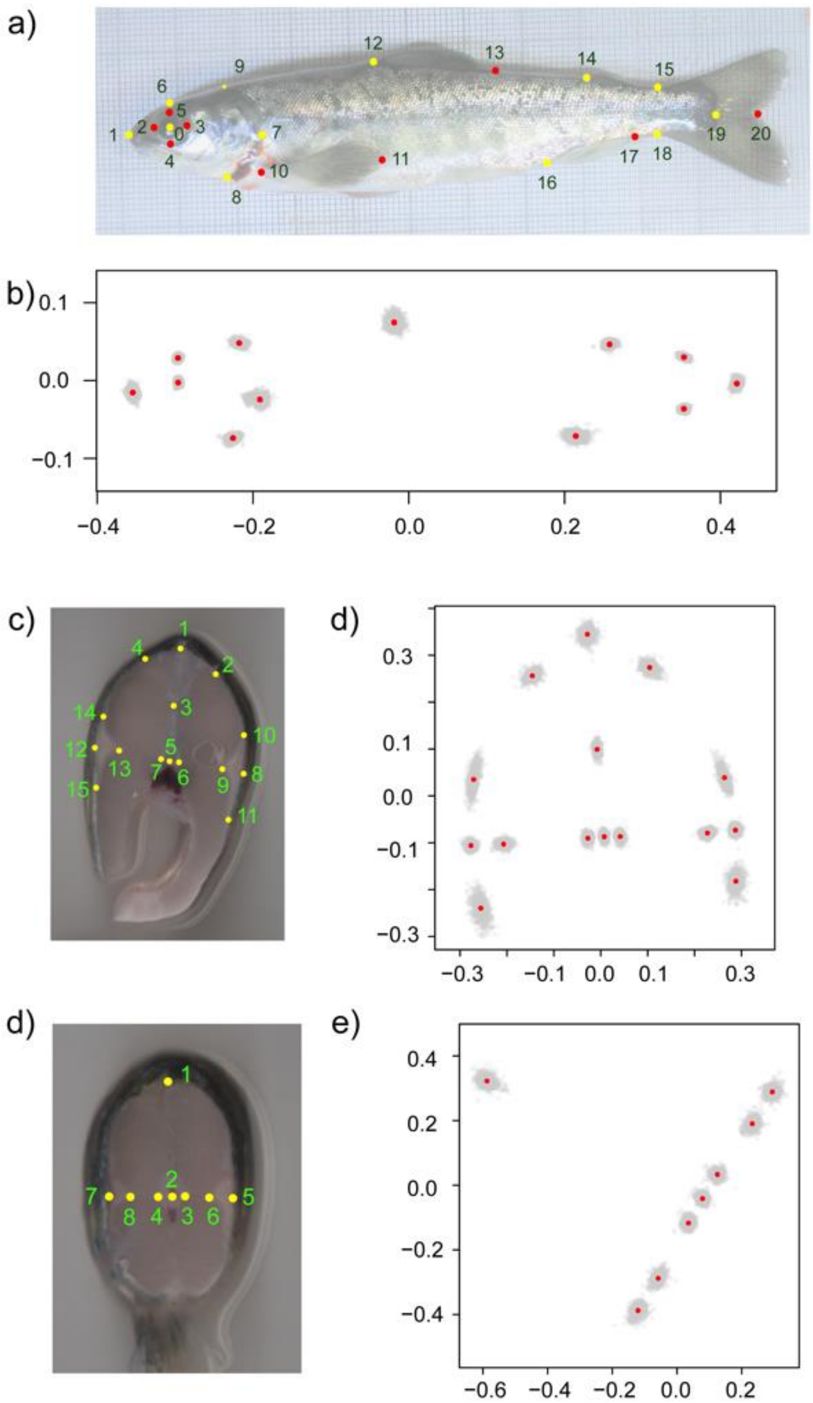


**Supplementary Figure 2**: Positions of landmarks (a, c, d), and coordinates of the Procrustes standardized landmarks, with red dots showing the mean position of the Procrustes standardized positions (b, d, e). Landmarks colored with yellow in (a, c, d) were used in the Procrustes analysis, while landmarks in red were used to calculate trait values (see Table 1 for the definitions of trait values). Axes indicate x- and y- coordinates of the Procrustes standardized landmarks.

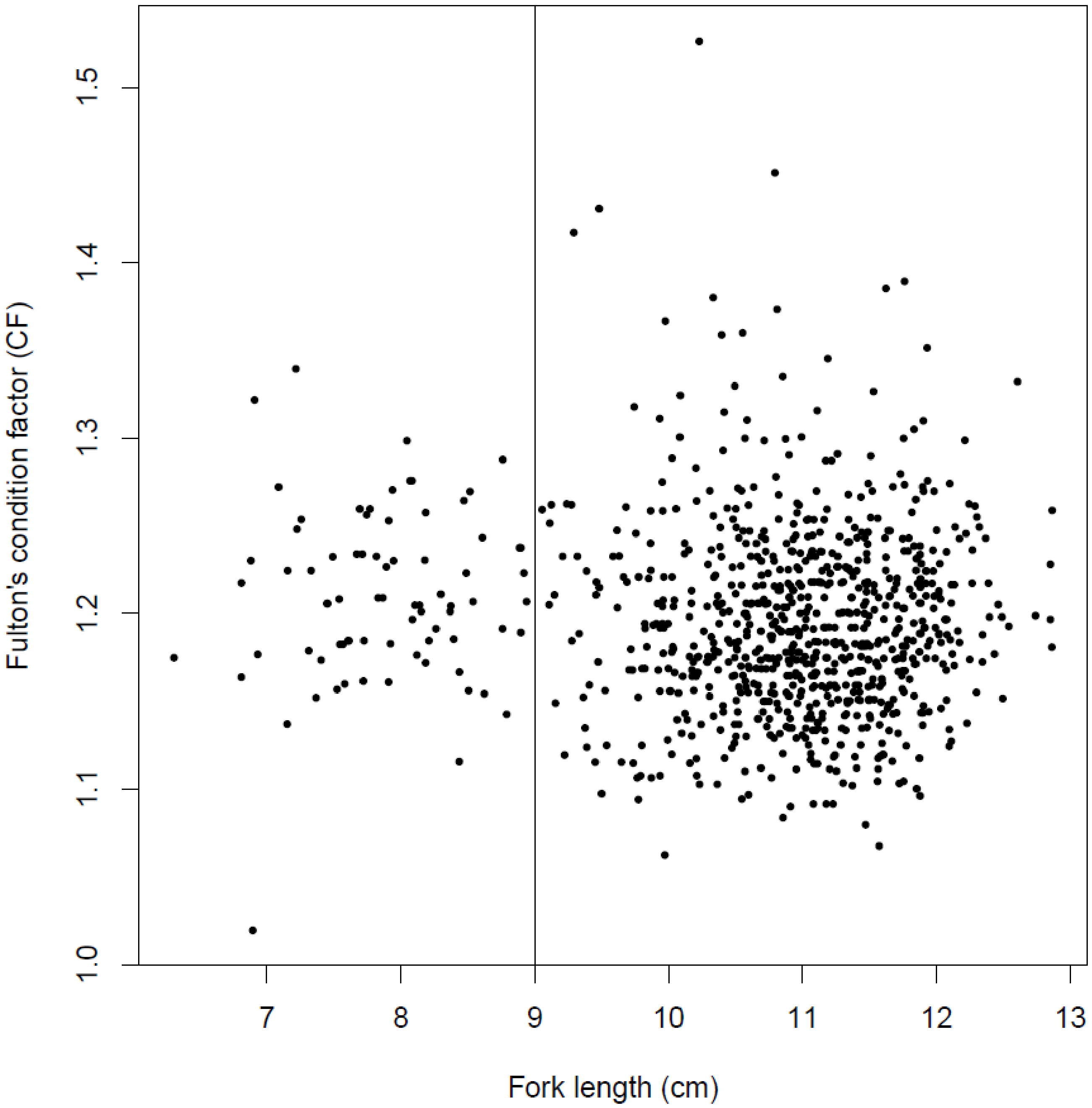


**Supplementary Figure 3**: Fork length plotted against Fulton's condition factor of the sampled fish. The horizontal line indicates the cut-off value at a length of 9 cm, that was used to filter fish grouped in the lower mode (N=69) prior to the analysis.

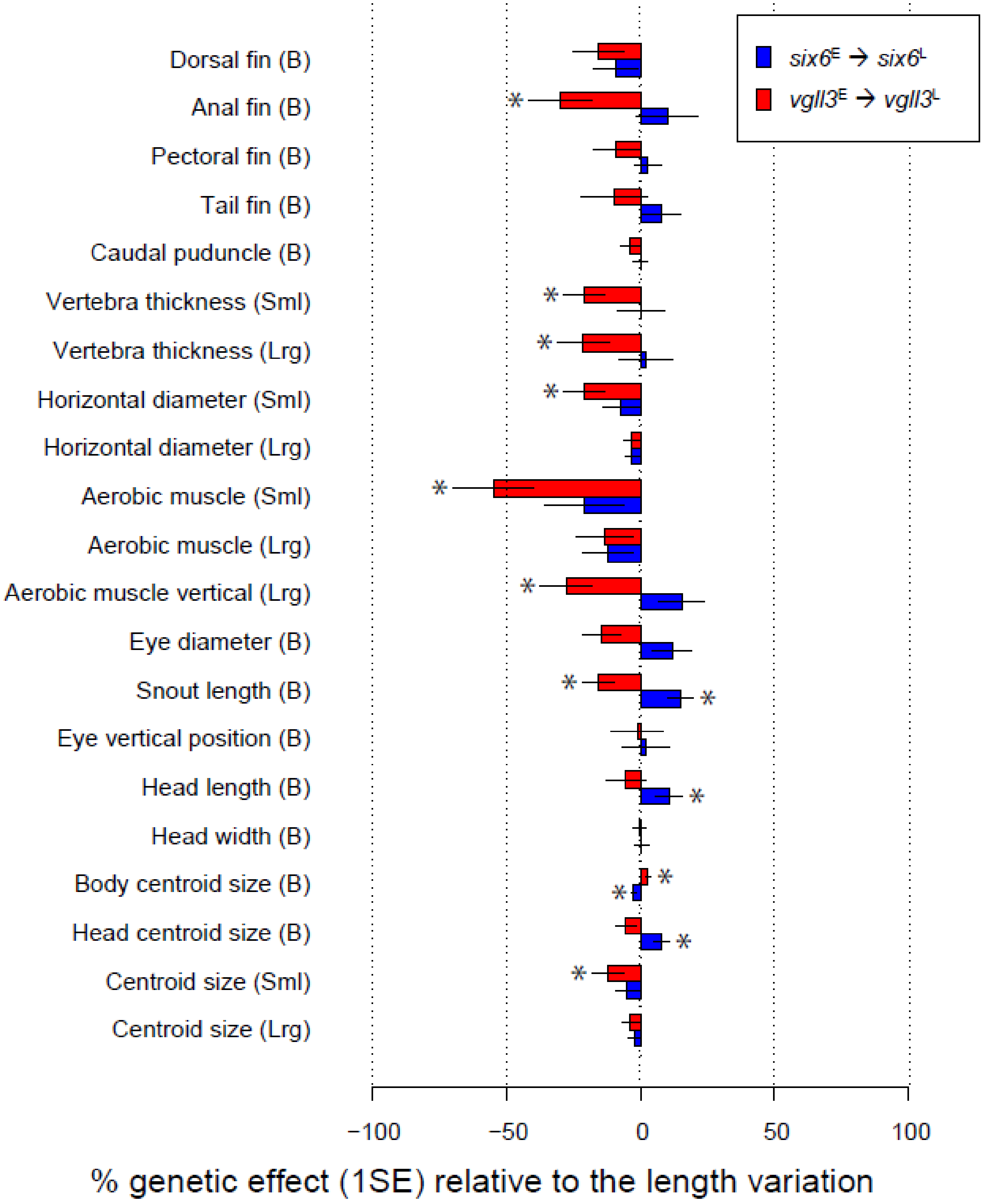


**Supplementary Figure 4:** Percent effect size difference between *vgll3*$^{EE}$ and *vgll3*$^{LL}$ (red lines), and *six6*$^{EE}$ and *six6*$^{LL}$ (blue lines) genotypes on trait values, relative to the effect of variation in total length (1 SD). Asterisks denote a significant genetic effect.

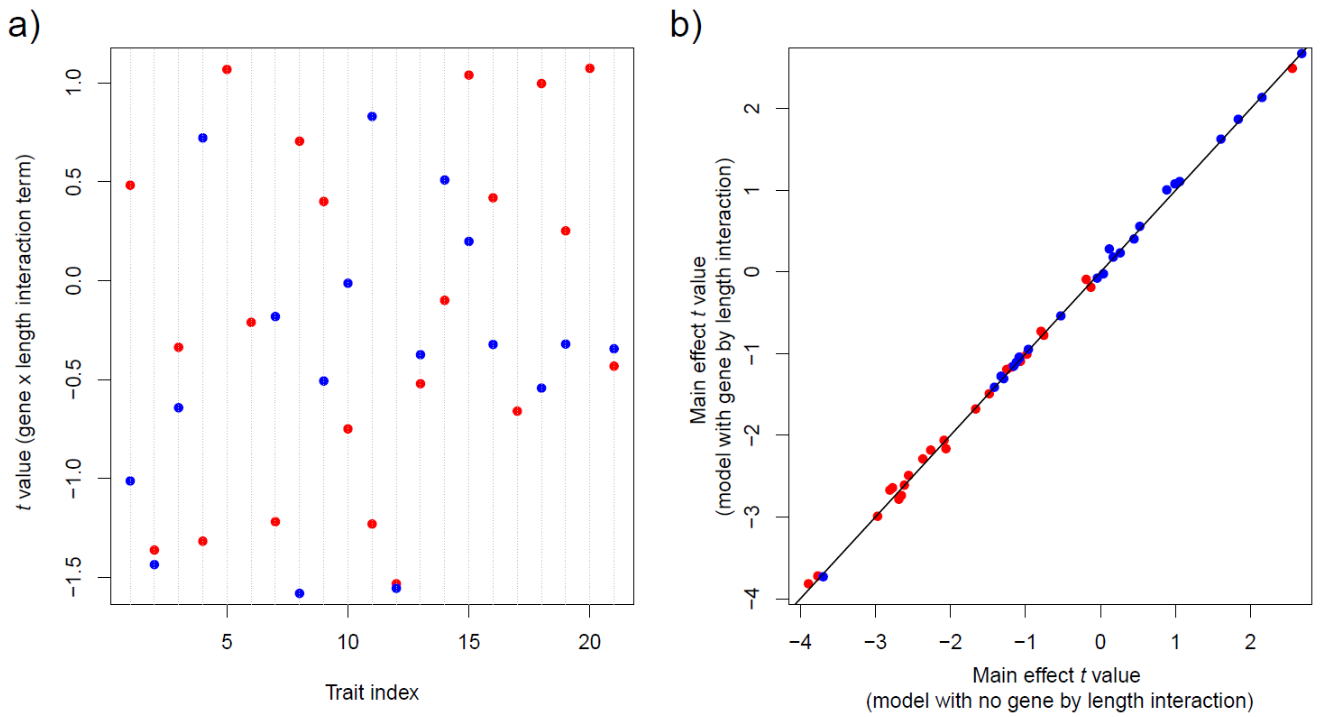


**Supplementary Figure 5:** The effect of including the gene by length interaction in models across 21 morphological traits. a) *t* values of *vgll3* x length (red) and *six6* x length (red) interaction terms in the interaction model. Traits are presented in the same order as in Table 1. b) The contrast of the main effect *t* value for vgll3 (red) and six6 (blue) terms between the main model (no gene by length interaction) and the interaction model. No change (y=x) is indicated by the black line.

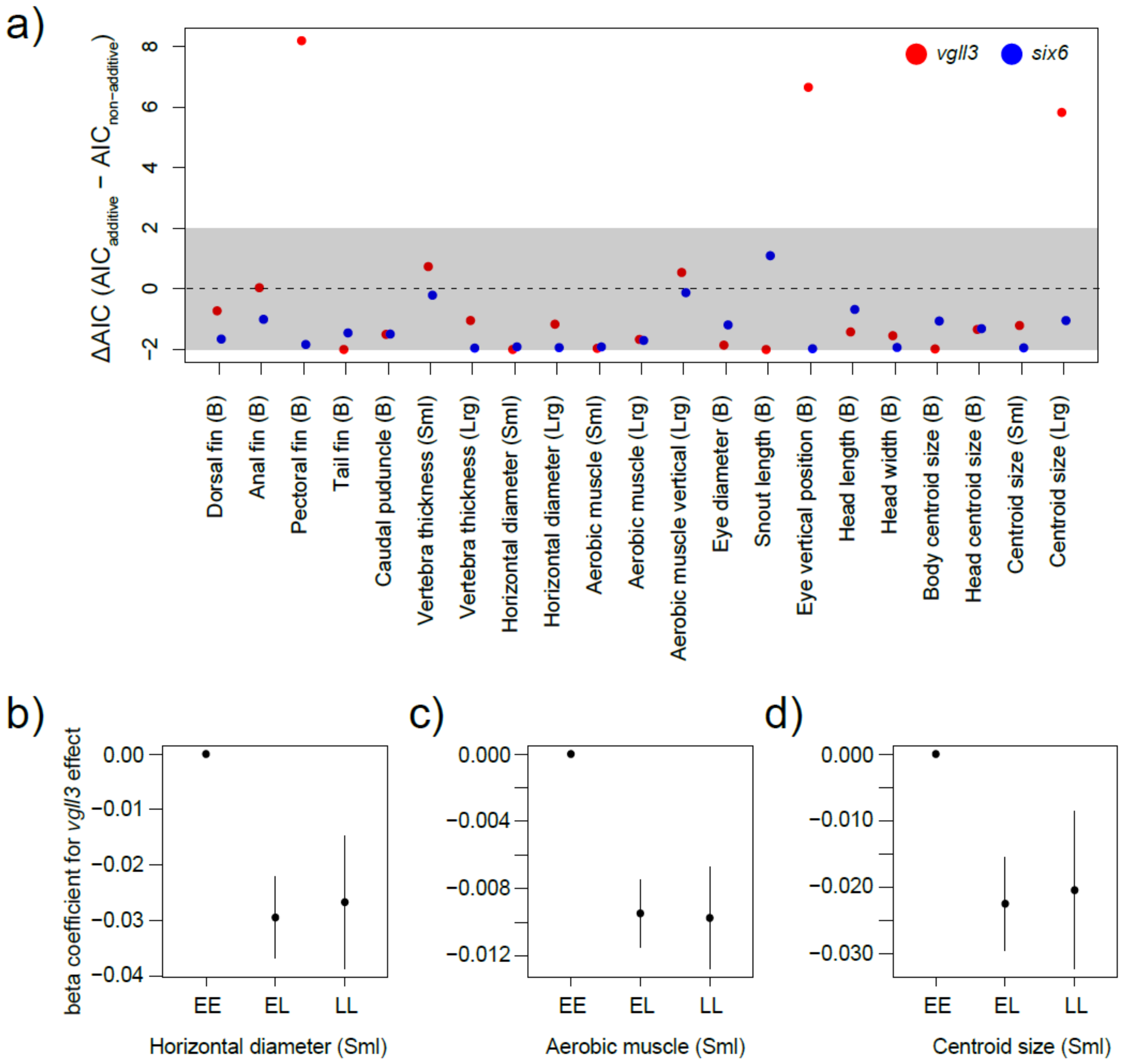


**Supplementary Figure 6:** a) Change in model parsimony when *vgll3* and *six6* effects were modelled non additively, across 21 traits. Gray shaded area indicates similar model parsimonies compared to additive model (i.e., ΔAIC ± 2 in non-additive model relative to the additive model). b-d) Trait genetic architecture of traits that exhibit improved parsimony (ΔAIC > 2) when genetic effect was modelled non-additively. Error bars indicate one standard error of the coefficient estimates of the model (relative to EE genotype effect).
